# Genome-wide association mapping and systems-level analysis reveal genetic architecture and physiological mechanisms linked with tolerance to flooding during germination in rice

**DOI:** 10.1101/2021.05.09.443312

**Authors:** Frederickson D. Entila, Myrish A. Pacleb, Evangelina S. Ella, Abdelbagi M. Ismail

## Abstract

Rice is the staple food of more than half of the world’s population; yet, it faces numerous challenges to meet the rising food demands and worsening climates. An urgent global imperative is to address imminent food shortages through intensive and sustainable agri-food systems and steady genetic gains. Intensification of rice production through direct-seeded rice (DSR) has been progressively practiced but is hindered by poor germination of existing high-yielding varieties in flooded soils. Identifying donors of anaerobic germination (AG) tolerance in rice will expedite the development of varieties suitable for DSR and will lead to enhanced and sustained agricultural productivity. This study aims to dissect the genetic architecture and physiological mechanisms of AG tolerance using systems biology and omics approaches. A Rice Diversity Panel (343 accessions) consisting of 5 subpopulations was screened for AG tolerance under greenhouse conditions, mapped through genome-wide association study (GWAS), and profiled for metabolites. Analyses revealed that most of the AG-tolerant varieties are japonicas with few indicas) and aus. Tolerant japonicas employed better root growth or rapid shoot extension, while tolerant indicas exhibited only the latter. A total of 51 significant GWAS peaks were detected across the genome, some of which were co-localized with known quantitative trait loci while others were novel, more so tolerance was found to involve different genetic controls across subpopulations. AG stress causes distinct biochemical signatures for tolerant genotypes and the profiles contrast among subpopulations implicating divergent metabolic adjustments, including shifts in sugars, intermediates, amino acids, antioxidants, and hormones. This study provides a systems-level approach for underpinning physiological mechanisms of AG tolerance; elucidating phenotypic heterogeneity, genetic architecture, transcriptomic networks, and metabolic landscapes from a genome-wide perspective.

**ONE SENTENCE SUMMARY:** The integration of GWA mapping, gene network analysis and, non-targeted metabolite profiling elucidates genetic architecture and physiological mechanisms of tolerance to germination and early seedling growth under anaerobic conditions in rice.

## INTRODUCTION

Arable land for food production is globally declining and climate is unfavorably changing, with natural resources continually deteriorating, while the demand for food of dramatically bloating population is rising. Furthermore, food production is being challenged by increasing urbanization and negative impact of climate change through loss of land and productivity, and decline in labor forces for agricultural purposes (<u>Masters et al. 2013</u>). Apparently, regions of the world that will be severely inflicted by climate change coincide with economically struggling countries, wherein hunger and undernourishment has been prevalent, and subsistence is primarily dependent on agriculture, thus posing more risks of food insecurity and aggravating socio-economic conditions (<u>Wheeler and von Braun, 2013</u>; <u>Morton, 2007</u>; <u>Howden et al. 2007</u>; <u>Hirabayashi et al. 2013</u>, <u>Schmidhuber and Tubiello, 2007</u>). These emerging problems substantiate the urgent need to enhance adaptation of agricultural crops to these adversities and through intensification of cultivation, sustainable food systems, and augmented genetic gains (<u>Howden et al. 2007</u>, <u>Godfray et al. 2010</u>; <u>Foley et al. 2011</u>).

Rice, being the staple food of more than half of the world’s population, faces numerous challenges to meet the rising future demands. To intensify rice production, reduce the costs and drudgery on farming communities, direct seeded rice (DSR) is being increasingly adopted in irrigated and rainfed areas as opposed to traditional transplanting. DSR involves establishment of rice from seeds directly sown in the field (<u>Farooq et al. 2010</u>). The shift towards DSR has several benefits: less labor requirement, cost- effective, simple, compatible for mechanization, leads to earlier maturity and less methane emissions, and reduce water requirements (<u>Sasaki, 1974</u>; <u>Balasubramanian and Hill, 2002</u>; <u>Pandey and Velasco,</u> <u>1999</u>). However, weed competition remains a major constraint in its wide implementation; hence farmers heavily rely on herbicides to manage weed infestation (<u>Tuong et al, 2000</u>). Though imposing early flooding could effectively control weeds in a sustainable manner, existing rice varieties can not germinate and establish in flooded soils, leading to failure of seedling establishment, especially if the field is poorly leveled or heavy rainfall occurs after seeding. As a result, farmers tend to use higher seed rates for DSR to compensate for early mortality and to ensure reasonable crop establishment, but at increasing costs (<u>Farooq et al. 2010</u>). Improved varieties with enhanced tolerance of flooding during germination (anaerobic germination [AG]) are required for direct-seeded rice systems in both rain-fed and irrigated areas, to ensure good crop establishment and for weed control in intensive irrigated systems (Ismail et al., 2012).

Rice is the only cereal that can germinate under hypoxic conditions, but this is limited to coleoptile extension with concomitant impedance of radicle protrusion (<u>Taylor, 1942</u>; <u>Ella and Setter, 1999</u>; <u>Lasanthi-Kudahettige et al. 2007</u>). The adaptation of rice to a wide range of hydrological environments allows the exploitation of existing plasticity and diversity for crop improvement through genetic manipulation (Ismail and Mackill, 2014). Genetic variation in rice responses to wet DSR has been previously reported (<u>Yamauchi et al. 1993</u>) and a number of donors for AG tolerance has been identified, which has recently been used in bi-parental mapping to identify quantitative trait loci (QTL) suitable for breeding programs. These studies facilitated the dissection of possible physiological machinery and discovery of traits tightly linked to the tolerance mechanism (<u>Ismail et al. 2009</u>; <u>Angaji et al. 2010</u>; <u>Septiningsih, et al. 2013</u>; <u>Baltazar et al. 2014; 2019</u>). Among cereals, rice has the enzymes required for starch hydrolysis and utilization of sugars under anoxia. Most rice genotypes are capable of initiating germination but failed to elongate further, only tolerant varieties were able to emerge due to efficiency in utilizing reserves under oxygen limitation (Miro and Ismail, 2013; 2018). Under low oxygen stress, sugar starvation and oxygen deprivation coerces Ca^2+^ release from mitochondria. The signal ions activate calcineurin B-like (CBL) protein forming CBL/Ca^2+^ complex which interacts with CBL-interacting protein kinase 15A (CIPK15A) that subsequently up-regulates SNF1-related protein kinase 1A (SnRK1A). The stimulated SnRK1A induces the promoter of transcription factor MYBS1 and also phosphorylates it to its active form. Activated MYBS1 protein binds to sugar response element (SRE) promoter forming MYB-DNA complex that substantially activates expression of α-amylases, particularly Amy3a subfamily (<u>Lee et</u> <u>al. 2009</u>; <u>Lu et al. 2007</u>; <u>Hong et al. 2012</u>; <u>Park et al. 2010</u>; <u>Loreti et al. 2007</u>). Subsequently, anoxia-induced amylases mobilize the starch reserves to supplicate energy and anabolic requirements of elongating embryonic axis (<u>Lee et al. 2014</u>). Metabolically, as repercussion of deficient oxygen supply, a dramatic shift from aerobic respiration to anaerobic fermentation commences. Despite its inefficiency, energy generation through substrate-level phosphorylation would suffice to address ATP crisis, thus increased rates of fermentative pathway remains integral for AG tolerance. Enzymes involved in alcoholic fermentation: pyruvate decarboxylase (PDC), alcohol dehydrogenase (ADH), and aldehyde dehydrogenase (ALDH), are substantially up-regulated under anaerobic germination (Ismail et al., 2009).

Recently, the genetic determinant for the major QTL for AG tolerance (AG1) derived from the rice landrace Khao Hlan On (Angaji et al., 2010), was identified as trehalose phosphate phosphatase, OsTPP7, involved in trehalose-6-phosphate (T6P) metabolism. OsTPP7 activity significantly promotes sink formation through superficially declining sugar availability by elevated T6P turnover. As a metabolic consequence, starch mobilization is enhanced and sustained, thus providing energy for coleoptile elongation, ultimately conferring AG tolerance (<u>Kretzschmar et al. 2015</u>). Though significant progress has been made in elucidating the metabolic adjustments and signaling cascades of germination under low oxygen conditions, further work is still needed to underpin the regulatory, signaling and physiological apparatus for AG tolerance due to the complex nature of the trait. Moreover, previous studies are limited only to a few genotypes not encompassing the breadth of natural variations and functional diversity that exist in rice gene pool.

Genome-wide association study (GWAS) has been progressively used to dissect the genetic architecture of complex traits in rice such as agronomic attributes, root and leaf morphology, metabolite profile, flowering time, tolerance to ozone, aluminum and salinity tolerance, and blast resistance (<u>Huang et al.</u> <u>2010</u>; <u>Yang et al. 2010</u>; <u>Ueda et al. 2015</u>; <u>Kumar et al. 2015</u>; <u>Courtois et al. 2013</u>; <u>Matsuda et al. 2015</u>; <u>Famoso et al. 2011</u>; <u>Wang et al. 2014</u>, <u>Huang et al. 2011</u>). GWAS provides a promising platform for linking phenotype and genotype to ascertain genomic regions explaining the trait of interest using diverse set of germplasm. Unlike QTL analysis, which is limited to recombination events from fixed generation following bi-parental crosses, GWAS utilizes variation existing in the diverse natural populations capturing wider allelic pools due to natural selection and domestication pressure that has occurred throughout the evolutionary course of the crop, thus leading also to high mapping resolution of genomic regions associated with the trait. Currently, the genomic resources for crops are dramatically increasing due to the plummeting costs for sequencing; hence genetic information will be more accessible, facilitating GWAS advancement and wide-spread use. The GWAS approach would enable identification of donors for a particular trait in a facile and manageable approach, hence expediting crop improvement and facilitating varietal development. This approach could also unearth the complex genetic architecture of the natural variations present in rice thus facilitates the elucidation of the physiological mechanisms associated with the phenotypic response. However, GWAS results are only limited to detecting genomic regions linked to the trait of interest but not directly pinpointing particular genes. Moreover, complex phenotypes or traits with low heritability as explained by many genetic elements of small effects with weak but genuine associations suffer from stringent significance thresholds for GWAS correction and were often neglected. These lapses are addressed through complementary approaches such as network analysis and pathway enrichments. Integrating GWAS results, transcriptomic network depictions, and pathway analysis could shed light on the systems-level functional mechanisms behind complex phenotypes and confidently identify genetic elements conferring the trait.

This study attempts to: (i) dissect the genetic architecture of tolerance to flooding during germination in rice using a diverse panel largely representing entire rice subpopulations; (ii) determine genomic regions highly associated with the phenotypic response; (iii) integrate systems-biology approach to uncover biological processes and molecular functions involved in the response; (iv) identify highly plausible genetic elements underlying the response; and (v) discover metabolic adjustments employed by tolerant genotypes from different subpopulations.

## RESULTS

### Phenotypic diversity reflects genetic heterogeneity

The Rice Diversity Panel 1 consisted of 343 inbred accessions with representatives across different subpopulations of *Oryza sativa* and collected from 82 countries. This panel was phenotyped for AG tolerance under greenhouse conditions and evaluated for 47 directly measured or derived traits under AG stress using two screening protocols. The traits include survival, plant morphology, and biochemical attributes (see Supplementary Table S2). Seed trays were used in the first method and root trainers in the second to allow large volume of soil for root growth.

Significant differences in tolerance (in comparison with tolerant Mazhan Red and sensitive IR42) among subpopulations were observed under AG conditions (p = < 0.001, Figure 1A). Most of the tolerant genotypes belong to the temperate and tropical japonica subgroups (means of 42.5% and 24.6%, respectively at 14 DAS), with fewer representatives from indica and aus, and with most of the aromatic subgroup being sensitive to AG stress (7.6% emergence at 14 DAS). Hierarchical clustering revealed tolerant clades, mostly dominated by temperate and tropical japonicas, with the tolerant indica and aus interspersed (Figure 1D). Geographical distribution of AG tolerance reveals variations across climatic zones (Figure 1E). Most of the tolerant accessions are located in temperate regions, coinciding with the origin of the temperate japonicas, which were mostly tolerant; roughly suggesting linkages of AG tolerance with tolerance of cold stress. Conversely, sensitive accessions are distributed more widely. Though phenotypic geographical distribution appeared to indirectly connect geo-climatic dependence of the responses, ecotypes from permanently and/or recurrently flooded conditions are still novel sources of tolerance.

**Figure 1.**
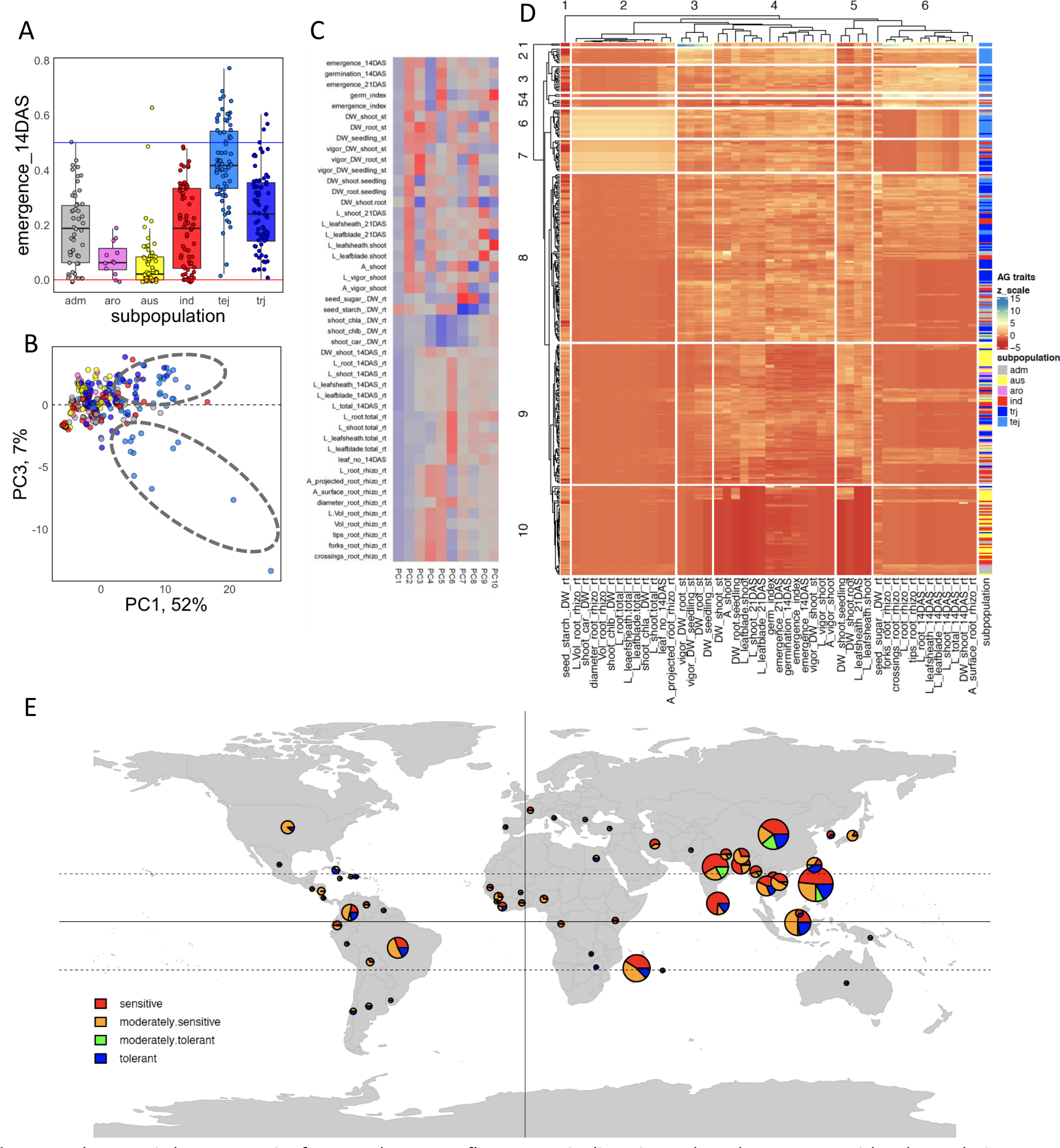
Phenotypic heterogeneity for AG tolerance reflects genetic diversity and tends to concur with subpopulation structure and geographical origin. Distribution of % emergence 14 DAS for each subgroup: adm=admixed; aro=aromatics; aus; ind=indica; tej=temperate japonica; trj=tropical japonica; blue line indicates performance of the tolerant check MaZhan Red and red line for the sensitive check IR42 (A). Phenotypic principal component (PC) analysis showed overlapping responses but tolerant japonicas and indicas deploy varying growth behaviors under AG stress (B-C). Heatmap and hierarchical clustering of the accessions using the phenotypes, color of row-side bars indicate subpopulations (D). Geographical distribution of tolerance revealed sporadic spread of sensitive genotypes while tolerant genotypes appear to coincide with the temperate zones (E).

Based on phenotypic principal component (PC) analysis, PC 1 to PC 4 could explain 84.8% of the phenotypic variations (Figure 1B). The pPC1 (51.9% of phenotypic variances explained) represents most of the traits linked to tolerance while pPC3 (6.7% of the phenotypic variances explained) scored higher eigenvector values for shoot related traits, but lower scores for root phenotypes (Figure 1C). Plot of PC1 and PC3 (Figure 1B) showed ambiguous clustering among subpopulations suggesting overlapping responses under AG stress; notably tolerant genotypes veer away from the overlap. Tolerant japonicas have either rapid shoot elongation or better root growth (high or low PC3); while tolerance of indicas is associated only with high shoot elongation (high PC3). These findings suggest possibilities for different tolerance strategies among subpopulations or evolution disparity due to selection sweep in relation to the source allocation and differential tissue growth for survival under AG conditions. Pearson correlation coefficients among measured and derived phenotypes were positive with the exception for seed starch (Suppl. Figure S1); likely due to mobilization of starch to soluble sugars to meet the metabolic demands of the growing embryo.

### Genome-wide association mapping links AG traits to numerous loci

Since ecological adaptation and subpopulation structure tends to concur with AG tolerance, as suggested by geo-climatic distribution (Figure 1E), subpopulation-specific responses, and the complexity of the trait, GWA mapping was accomplished for individual subpopulations besides the whole panel. Due to small population sizes for stratified association study, closely related subgroups were merged to form reasonable sample sizes: indica and aus to form ind_aus panel (n=128); temperate japonicas, tropical japonica and aromatics to form jap_aro panel (n=158); and lastly the whole panel with all accessions.

GWAS was implemented in ECMLM algorithm and the first two PCs and kinship estimates were included as covariates to correct for population structure for both whole set and stratified analysis. Most of the SNP peaks from the analyses did not reach the stringent Bonferroni threshold (<u>Holm, 1979</u>), though distinct associations were observed. To improve the statistical power of association mapping, the SUPER algorithm was implemented in the analysis. This method dramatically reduces the number of SNPs to be tested in determining genetic relationships, thus considerably increases statistical power. A total of 39, 11, and 1 SNPs surpassed the stringent Bonferroni threshold (-log10 p-value > 5.81) and were considered significant associations from the all, ind_aus, and jap_aro panels, respectively (Suppl. Table S3). The significant SNPs were associated with leaf sheath length, shoot length, and shoot dry weight for the all panel; root diameter, seed starch, total length, root volume, and leaf blade length for ind_aus panel; and root dry weight for jap_aro panel. Some of the significant SNPs co-localized or are in close proximity with published QTLs linked with AG tolerance (Figure 2, Suppl. Figures S2-S7) particularly in chromosomes 1, 7, 8 and 9. Notably, some of the significant SNPs detected novel associated regions in the rice genome, not reported before. Interestingly, when considering the SNPs with –log 10(p-values) > 4 from all of the traits, population-specific alleles become virtually undetected when the whole panel is analyzed; few associated SNPs were shared between panels (around 3-8 SNPs) and remarkably only one SNP in chromosome 4 (id4000981) has been detected to be commonly shared among the all, ind_aus, and jap_aro panels (Figure 2B). As summarized in Figure 2A (see also Suppl. Figures S2-S7 for details), the genetic architecture of AG traits differs significantly among the subpopulations and different GWAS peaks were detected when the analysis is conducted individually on merged subpopulations or when the whole diversity panel is analyzed. This strongly indicates that allelic variations linked with AG tolerance across subpopulations occur in varying genomic addresses. Distribution of MAF indicates that significant allelic variations are quite less frequently found in the diversity panel (Suppl. Figure S8), with average MAF of 0.23 - 0.28 and median MAF of 0.22- 0.30 for the panels. A number of significant SNPs were also found to be linked with more than one trait, implying pleiotropic effects of the allelic variants. However, these alleles tend to appear in lower frequencies in the rice population being studied. Notably, heritability values were also low (0–19.8%, 0.4–29.5% and, 0–19.4% for all, ind_aus and, jap_aro panels, respectively). Association hot spots are conspicuous in chromosome 1 for most of the panels, likely because most of the abiotic tolerance genes are situated in this portion of the rice genome. Moreover, associated subpopulation specific variations were concentrated in certain regions as well; chromosome 3, 9, and 11 for ind_aus panel, and chromosomes 8 and 11 for jap_aro panel (Figure 2A).

**Figure 2.**
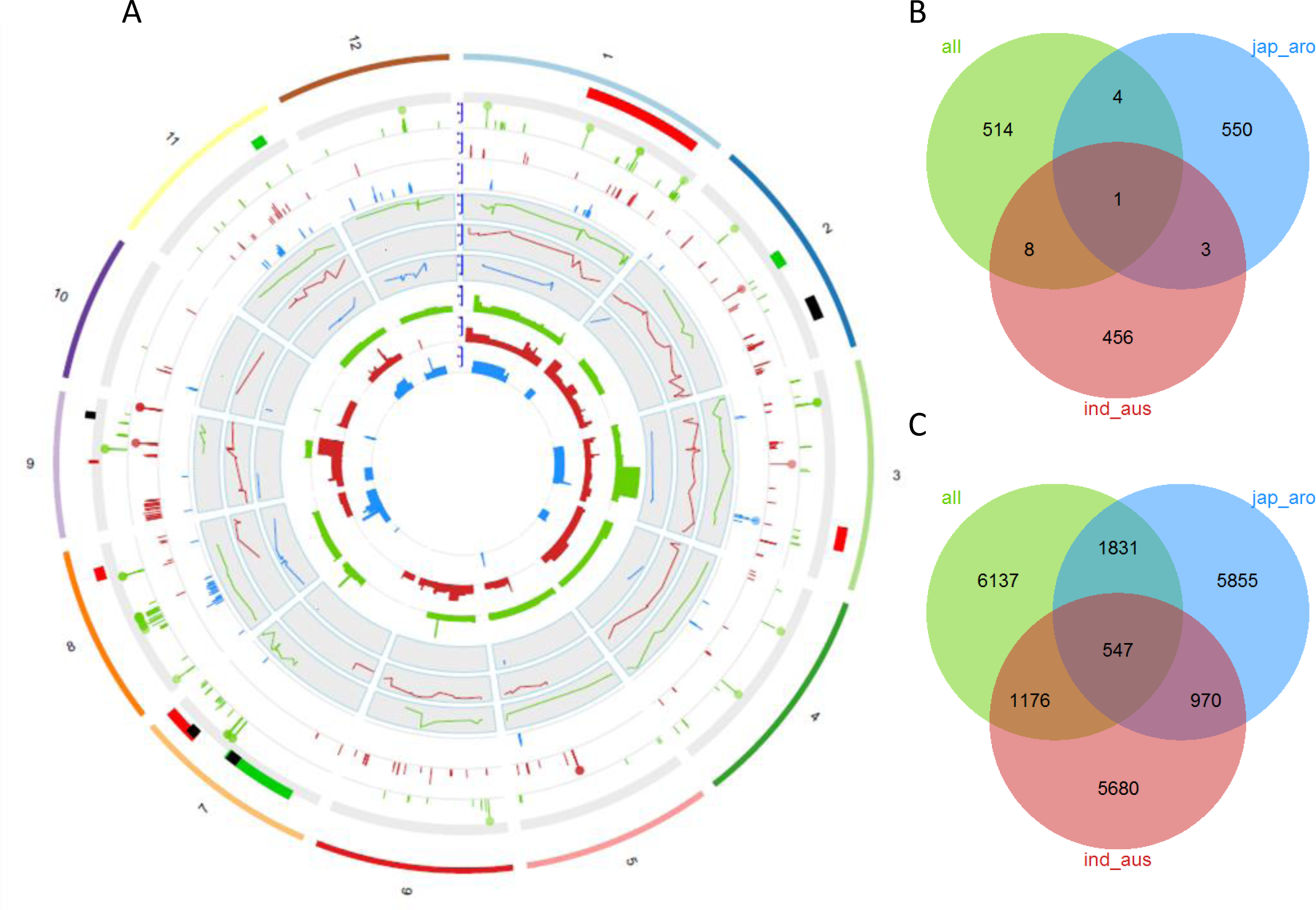
Genomic regions associated with AG traits: first track includes published QTLs linked with AG tolerance derived from different donors, dark red (KHO), dark green (Nanhi), black (Mazhan Red); second to fourth track indicate position of top 20 SNPs from each trait, with the bar representing –log10 p-values, filled circles identify significant associations, for the all (green), ind_aus (red), and jap_aro (blue) panels. Fifth to seventh track include line graphs for minor allele frequency from ECMLM approach; eighth to eleventh track shows number of traits associated with the SNPs (A). Venn diagram of overlapping SNPs associated with traits for the stratified analyses, including the all, ind_aus and, jap_aro panels (B). Venn diagram of overlapping candidate genes inspected from ± 100kb of the SNPs (C).

### Conventional pathway enrichment uncovers biological processes involved in AG response

To generate a list of initial candidate genes from the association mapping, the top 20 SNPs of each trait were further inspected. SNPs were merged within 200kb span (± 100kb from the SNP position) to form LD blocks. This genomic window was selected since linkage disequilibrium tends to decay beyond this distance. This modest window is also close to the average LD decay of different rice subpopulations: ≈100kb indica, 200kb in aus and temperate japonica, and 300 kb in tropical japonica (<u>Zhao et al. 2009</u>). Combining the LD blocks and singletons, there were a total of 252, 213, and 238 regions for the all, ind_aus, and jap_aro panels, respectively with which one SNP is common among panels (Figure 2B). Upon close inspection of these regions, a total of 9691, 8373, and 9203 gene models were considered putative determinants of the trait from the all, ind_aus and jap_aro panels, respectively (Figure 2C). When the entire regions of 200kb span is considered, 547 gene models interspersed in entire rice genome except chromosome 12, were found to be common among the panels, suggesting that these genetic elements may play central role in global response to AG stress across subpopulations.

These gene lists were examined for enrichment or discrimination for certain functional themes: biological processes, molecular function, cellular component, and protein class, through canonical pathway enrichment analysis using the PANTHER database (<u>Mi et al. 2013</u>). For the all panel, a total of 14 and 5 GO terms for biological processes category were significantly enriched and discriminated, respectively. The most significant was protein phosphorylation (GO: 0006468) with 2.70** fold enrichment. Other significant terms included cell surface receptor signaling (GO: 0007166), response to stress (GO: 0006950), and organelle organization (GO: 0006996) with corresponding fold scores of 2.47**, 1.53*, and -0.21** respectively. For the ind_aus panel, a total of 6 terms were significantly overrepresented and 4 GO terms were significantly underrepresented. These include protein phosphorylation (GO: 000646), mitosis (GO: 0007067) and carbohydrate metabolic process (GO: 0005975) with corresponding fold scores of 2.21*, 1.97** and, -0.58*. For the jap_aro panel, 14 GO terms were significantly enriched and 21 GO terms significantly discriminated. These comprised of signal transduction (GO: 0007165), primary metabolic process (GO: 0044238), and protein transport (GO: 0015031) with overrepresentation scores of 2.06***, -0.77** and, -0.43*** respectively. The 547 genes shared among panels were significantly involved with biological processes particularly metabolic process (GO: 0008152), and some with unclassified functions with fold enrichments of- 0.24** and, 1.16*, respectively. Though conventional pathway-based analysis revealed biological and molecular functions associated with the gene sets, the identified terms are general in nature and do not particularly identify definite mechanisms underlying variations in AG tolerance (Suppl. Dataset S2).

### Weighted gene co-expression network showed distinct modules highly pertinent to AG response

To generate gene co-expression network, we used microarray data of germinating seeds grown in normoxic and anoxic conditions of 16 replicated samples (<u>Lasanthi-Kudahettige et al. 2007</u>; <u>Narsai et al.</u> <u>2009</u>) with corresponding coleoptile measurements imputed. Expression profiles for the candidate genes were extracted from the dataset: 3234, 2572, and 2534 candidate genes generated for the all, ind_aus, and jap_aro panels (Figure 3A, Suppl. Figures S9A and S10A). The weighted gene co-expression network analysis (WGCNA) approach was implemented to discover modules of highly correlated genes and relate these expression profiles to its corresponding phenotypic responses. The resulting modules usually comprised of genes involved in the same biological pathways, and more likely to share regulatory factors.

**Figure 3.**
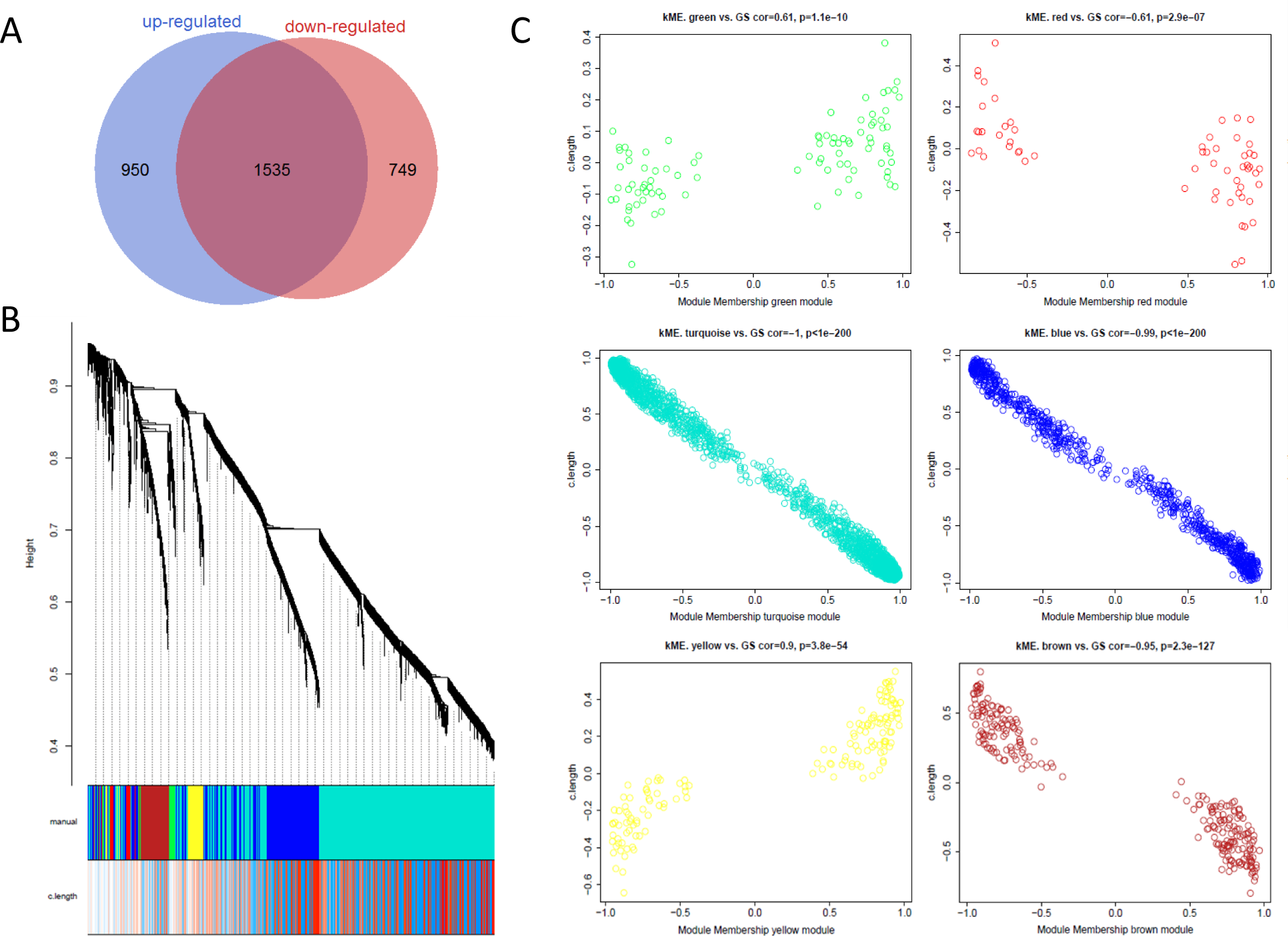
Venn diagram of differentially expressed genes for all panel (A). Co-expression network from the all panel, each line is an individual gene and the branches correspond to modules of highly interconnected genes, below the dendogram, each gene is color coded to indicate module assignment and the GS values for coleoptile length (B). Correlation between MM and GS for each of the identified GWAS modules, all of the modules had significant correlations (p-values < 0.01) for GS and MM (C).

Analysis of the all panel resulted in 6 distinct gene modules (Figure 3B, Suppl. Figure S18) with sizes ranging from 59 genes for the all red module to 1865 genes for the all turquoise module. These modules are large networks with highly connected genes (edge number extends 1658 to 1677028). Modules had considerable MS values ranging from 0.10 (all green module) to 0.68 (all turquoise module). A complete list of module assignments and network parameters for genes are included in Suppl. Dataset S1. Eigengene network showed that coleoptile length is highly related to the all yellow module. All of the modules had significant correlations for GS and MM (correlation values of -1.00*** to 0.90***) indicating that these modules harbor genes highly relevant to coleoptile length (Figure 3C). Analyses and significant modules for jap_aro and ind_aus sets are represented in Suppl. Figures S9, S10, S19, S20, and the complete list of gene module assignments is summarized in Suppl. Dataset 1.

To gain insight on the interaction of the genes across the genome in a global scale, the genes from the modules with strong correlation with coleoptile length (absolute GS values ≥ 0.80) were selected and the 97.5 percentile of the interactions were visualized (Suppl. Figure S11). Generally, only the large modules, turquoise and blue were preserved despite stringent thresholds, though few genes from brown and red modules for the all and jap_aro panels were included. Genes from significantly associated genomic regions are strongly co-expressed; likewise interact as well with genes in blocks of modest association which were below the stringent GWAS significance cut-off. This imply that highly plausible genes from genomic spans of true association but with small effects will be more likely to suffer when analyses rely solely on statistical evidences, which is the case for complex traits. Therefore, inclusion of regions of marginal significance is still substantial in elucidating trait physiology in the context of gene regulation and protein-protein interaction.

### Pathway-based analysis of module content revealed key specific roles for regulation of AG stress

Network analysis can extract more biological insights from GWAS by exposing specific biological mechanisms that are not observed when candidate gene list is analyzed entirely. Module content pathway analysis could uncover specific and even novel processes and functions condensed within modules (<u>Farber, 2013</u>). Remarkable enrichments were revealed upon module content characterizations of the all panel. The all blue module was found to be significantly involved in protein lipidation (GO: 0006497), carbohydrate transport (GO: 0008643), and protein metabolic processes (GO: 0019538) with fold enrichments of 21.4*, 9.8*, and 2.4* respectively. The all brown module seemed to play vital roles in response to endogenous stimulus (GO: 0009719), and tRNA aminoacylation for protein translation (GO: 0006418) with fold overrepresentation of 37.8*, and 37.1*, respectively. Genes from the all green module were engaged in DNA-dependent transcription (GO: 0006351), metabolism of nucleobase-containing compounds (GO: 0006139), and primary metabolism (GO: 0044238) with corresponding fold enrichments of 28.8**, 13.9***, and 5.5 **. The all red module had vital roles for apoptopic process (GO: 0006915), cell surface receptor signaling (GO: 0007166) and sulfur compound metabolism (GO: 0006790), fold overrepresentation of 66.9*, 40.1*, and 37.3*, respectively. The all turquoise module had extensive participation in cellular glucose homeostasis (GO: 0001678), response to abiotic stimulus (GO: 0009628), and protein phosphorylation (GO: 0006468), respectively with the following fold enrichments: 39.7*, 9.1*, and 8.8**.

Module content characterization of the ind_aus panel revealed various specific and novel pathways that canonical pathway-based analysis failed to pinpoint, particularly functions for metabolism, biological regulation and signaling apparatus. The ind_aus blue module had pathway categories significantly enriched for I-kappaB kinase/NF-kappaB cascade (GO: 0007249) and G-protein coupled receptor signaling pathway (GO: 0007186) with fold enrichment of 59.7*, and 23.6*, respectively. The ind_aus brown module is mostly involved in phospholipid metabolic process (GO: 0006644), carbohydrate transport (GO: 0008643), and tRNA metabolic process (GO: 0006399) with corresponding fold enrichments of 27.1*, 21.0*, and 20.7*. The ind_aus green module had condensed functions for receptor-mediated endocytosis (GO: 0006898), with fold overrepresentation of >100**. The ind_aus turquoise module genes participate in MAPK cascade (GO: 0000165), mRNA processing (GO: 0006397), and RNA metabolic process (GO: 0016070), with fold scores of 10.2**, 4.9**, and 3.7***, respectively. The ind_aus yellow module had functional overrepresentations for regulation of transcription from RNA polymerase II promoter (GO: 0006357) and signal transduction (GO: 0007165), with fold scores of 36.3* and 21.0* correspondingly.

Pathway-based analysis of the jap_aro panel uncovers specific functions tightly linked with central metabolism, regulation of biological processes, and signaling. Upon module characterization, the jap_aro blue module had overrepresented occupations for nitrogen utilization (GO: 0019740), and G-protein coupled receptor signaling pathway (GO: 0007186), with corresponding fold enrichments of >100**, and 30.5*. The jap_aro brown module had genes with overrepresented categories for mRNA polyadenylation (GO: 0006378), mRNA 3’-end processing (GO: 0031124), and mRNA splicing, via spliceosome (GO: 0000398), with fold increase of 46.0*, 41.6*, and 18.5**, respectively. The jap_aro turquoise module had term overrepresentation for cellular glucose homeostasis (GO: 0001678), amino acid transport (GO: 0006865), and polysaccharide metabolic process (GO: 0005976) with fold scores of 42.4*, 12.1**, and 5.0**, respectively. The jap_aro green module had extensive participation in signaling cascades: JNK cascade (GO: 0007254), transmembrane receptor protein serine/threonine kinase signaling (GO: 0007178), I-kappaB kinase/NF-kappaB cascade (GO: 0007249) and, MAPK cascade (GO: 0000165), with corresponding fold scores of > 100** for all terms. The complete list of enriched GO categories for the entire panel modules are summarized in Suppl. Dataset 2 and module network depictions in Suppl. Figures S18-S20.

### Allele mining revealed different putative gene candidates for AG tolerance among stratified analyses

From the genomic regions of significant associations, genes with absolute GS values ≥ 0.80 were selected as putative determinant of traits. Haplotype analysis was performed for the associated regions of the selected genes to determine haplotype blocks of strong linkage disequilibrium and identify allelic variants associated with AG tolerance; the highly connected neighboring genes were also assessed for functional condensation. In-silico analyses was extended on the selected genes by using the SNPs form the SNP-seek database extracted from 3000 rice genomes. Flanking sequences (2kb upstream and downstream of the gene) was also considered to capture variations in the regulatory elements. For the purpose of brevity, one gene from each stratified analysis with annotated functional roles of interest was chosen.

On the all turquoise module, LOC_Os08g34580 (GS=-0.82) is a putative trehalose-6-phosphate synthase with functional involvement in carbohydrate pathway; expression analysis revealed upregulation under anoxia (-logfc= 0.23***). The gene falls in the all LDb08009 (comprising of 3 significant SNPs: id8006298, id8006299 and, id8006308, Figure 4), which is tightly linked with leaf sheath length and shoot biomass. Inspection of the LD region revealed 2 blocks, with 2 haplotypes for the first block and 9 haplotypes for the second block. Distribution analysis showed that GCT allele strongly correlates with tolerance. Subnetwork examination revealed tight connections with 301 genes having roles in transcription regulation and carbohydrate metabolism (fold enrichments of 47.65*** and 7.31*, respectively), mostly glucosidases, kinase modulators and, G-protein coupled receptors. Inspection with SNPs from 3000 genomes revealed that variations coincide with subpopulation structure, with considerably low MAF (Suppl. Figure S14). Hierarchical clustering distinguishes tolerant phenotypes for temperate and tropical japonicas.

**Figure 4.**
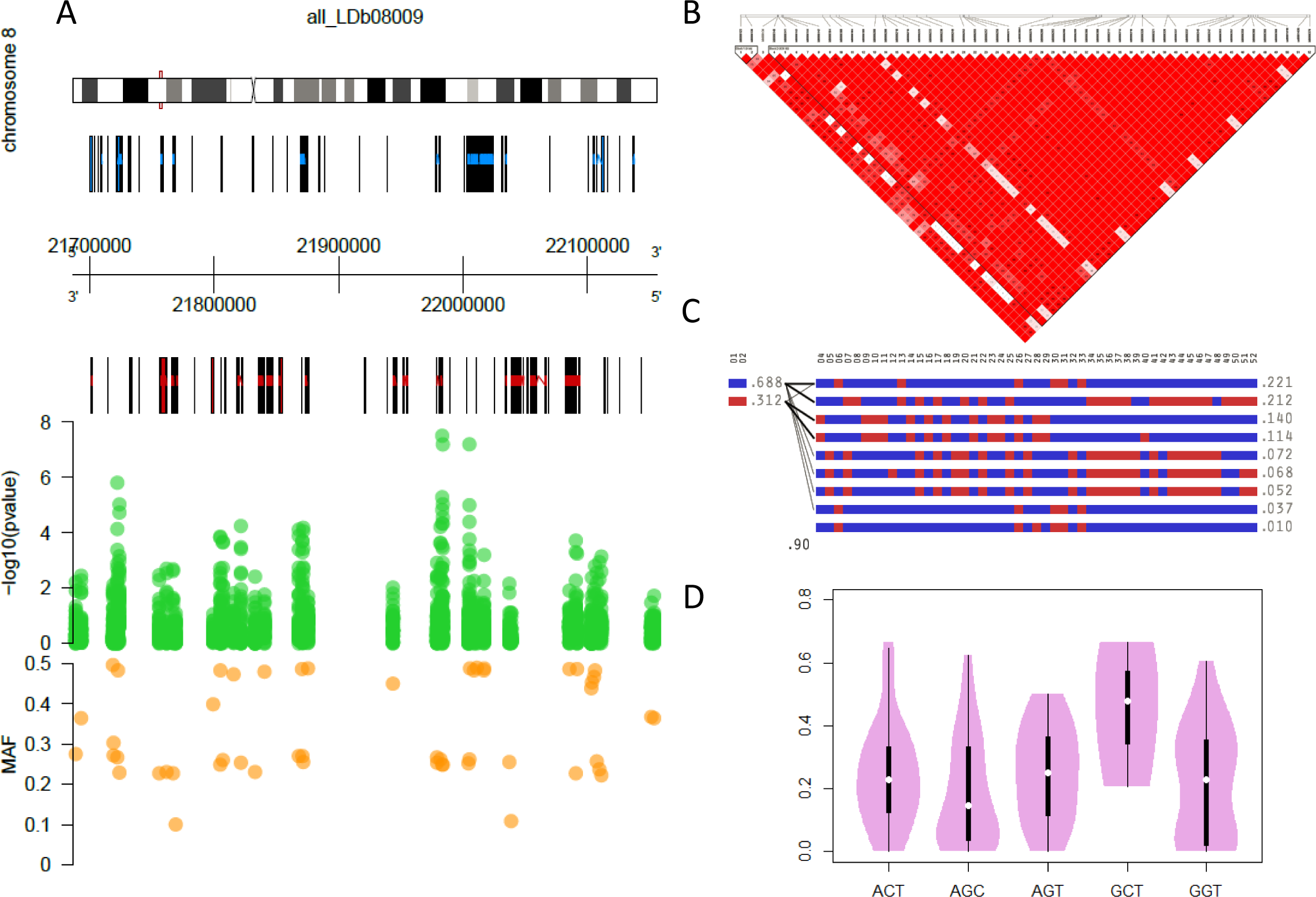
Ideogram depiction of genes present in all_LDb08009 region in chromosome 8 with corresponding GWAS significance values and MAF (A). Linkage disequilibrium heatmap of the region showing 2 haplotype blocks (B). Haplotype variants existing in the associated region with corresponding linkage disequilibrium value between blocks and the frequencies of the variants (C). Frequency distribution of the identified allelic variants and corresponding tolerance (D).

The gene LOC_Os09g26310 from the ind_aus turquoise module is a putative hypro1 glycosyl transferase functioning in carbohydrate metabolism and protein modification. This correlates strongly with coleoptile length and is repressed dramatically under anoxia (GS = 0.98, -logfc= -0.40***). The gene coincides with the ind_aus LDb09010 consisting of 2 significant SNPS: id9004677 and id9004707, strongly linked with shoot length, root length and leaf number (Suppl. Figure S12). The associated region consisted of 4 LD blocks with haplotypes ranging from 3 to 7 variants. Haplotype examination revealed that CCC allele is concomitant with AG tolerance but exist in small frequency in the ind_aus panel. It is highly co-expressed with 242 genes associated with mRNA transcription and protein methylation (fold enrichments of >100** and 31.2*, respectively). The variations in genomic region containing the gene correspond with subpopulation structure, however MAF are considerably high, in particular the coding and the 5’ UTR. The percent mismatch with the Nipponbare genome distinguishes tolerance in indicas (Suppl. Figure S15).

From the jap_aro blue module, LOC_Os03g50220, a putative protein kinase is highly associated with coleoptile growth and considerably down regulated under anoxic conditions (GS = 0.84, -logfc = - 0.14***). The gene falls within the LD block of the significant SNP id3013397 associated with vigor, root biomass and shoot traits (Suppl. Figure S13). Haplotype analysis revealed that the GCT allele concurred with tolerance, however exists in small frequency in the jap_aro panel. Dissection of the subnetwork with 341 genes uncovered pathway condensation for G-protein coupled receptor signaling (fold enrichment of 48.76*) and mainly interacts with other kinases and cysteine proteases. Clustering of the 3000 genome SNPs poorly separates the tolerant phenotypes. It can be assumed that the tolerance can be attributed to transcriptional and/or translational regulation of the gene, since it contains several exons that may form different splice variants (Suppl. Figure S16).

### Subpopulations employed distinct metabolic landscapes under AG stress

To understand early metabolic adjustments under AG perturbation, genotypes from different subpopulations with varying AG responses were selected for non-targeted metabolite profiling at 4 DAS, where tolerance phenotype is distinguishable. A mixture of different solvents was used as extractant to expand the diversity of compounds to assay, from polar to semi-polar for global purpose. Upon peak identification, 1355 metabolic features were detected with 166 identified metabolites. Analysis was done separately for ind_aus and jap_aro panels to determine subpopulation specific and core metabolic confluences under oxygen deprivation.

PCA analyses revealed that mPC1-mPC3 could explain 38.3% and 36.8% of the metabolic variations existing in jap_aro and ind_aus metabolomes, respectively (Figures 5B and F). Though no apparent separation for tolerant and sensitive genotypes was observed, mPC3 distinguishes tolerance under AG stress. To further reveal metabolic alterations, Bayesian statistics was conducted for all metabolic features. A total of 550 metabolic features were substantially up-regulated and 355 down-regulated as treatment effect in the ind_aus panel, while 579 metabolic features were significantly up-regulated and 295 significantly down-regulated as treatment effect for jap_aro panel. Most of the differentially fluxed metabolic features (714) were shared between panels implicating core metabolic shifts under AG stress. Significant increases in sugar and its derivatives were observed for both panels particularly glucose, glucose-6-phosphate, fructose and sucrose. Critical decline in fructose-6-phosphate (-logfc of -0.31* and -0.44** for ind_aus and jap_aro panels, correspondingly) but stable levels of downstream intermediate glycerate-3-phosphate may indicate increased glycolytic activity under AG stress. Variations in levels of TCA intermediates were observed between subpopulations (Figures 5K-L). For ind_aus panel, reduction in fumarate and maleate were remarkable (-logfc of -0.51** and -0.98***, respectively). However in jap_aro panel, only maleate decreased (-1.00***) while isocitrate and 2-oxoglutarate increased (-logfc of 0.82*** and 0.45**, respectively). This variation indicates that TCA cycle and probably glyoxylate shunt are considerably driven in jap_aro genotypes despite oxygen deficits for the purpose of recycling carbon sources. Likewise, the ind_aus genotypes might utilize the glyoxylate shunt to generate energy and provide substrates for protein biosynthesis. Accordingly, jap_aro panel had more diverse amino acids derived from TCA intermediates, particularly aspartate, asparagine and hydroxyproline, which were not elevated in ind_aus group. However aromatic amino acids tryptophan and tyrosine were copious for both panels. Accumulation of pyruvate occurred in both subpopulations as a repercussion of dampened TCA cycle. Consequently, oxygen deficit resulted in significant elevation of pyruvate products, lactate from fermentation pathways for energy production, and alanine for reconfiguration of N metabolism as resource economy. A number of alkaloids, particularly calystegine compounds were abundant under AG signifying ROS sequestration by these antioxidants. Some nucleotides were found to increase in both panels under AG stress, particularly guanosine, adenosine, thymidine, GMP and, UMP. The complete list of metabolic module assignments and enriched GO categories for the metabolic modules are summarized in Suppl. Datasets S3 and S4.

**Figure 5.**
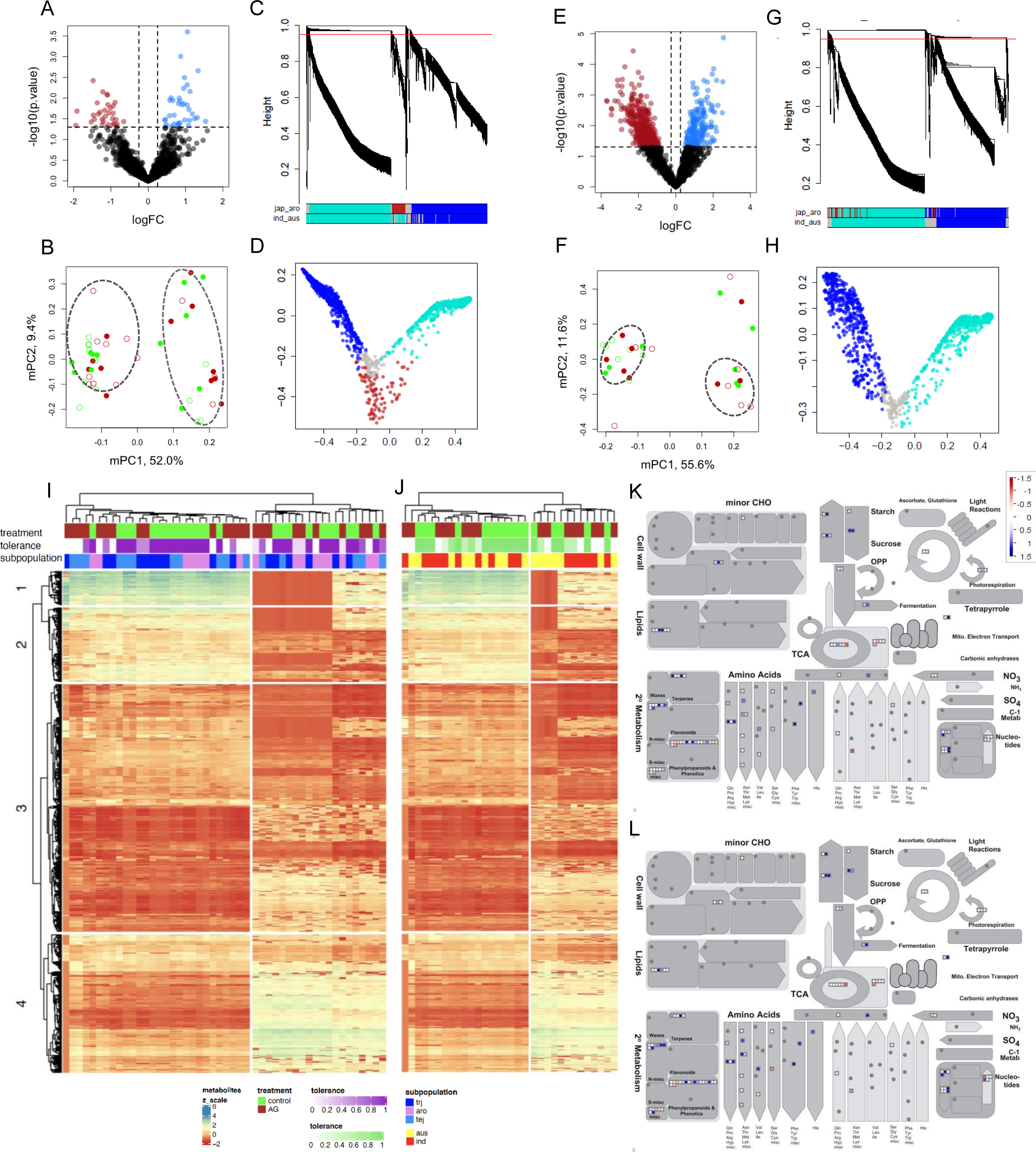
Metabolomic profiles of jap_aro (A-D, I, K) and ind_aus (E-H, J, L) panels of contrasting AG responses: volcano plots indicating differentially fluxed metabolic features for treatment*tolerance effects (A, E); metabolic Principal Component (mPC) analysis revealed distinct metabolic compositions under AG stress for representative tolerant (filled circles) and sensitive (unfilled circles) genotypes (B, F); Metabolic network structure and (D, H) multidimensional scaling of metabolite features showed highly conserved and distinct metabolic responses (C, G); heatmap representation of fluxed metabolic features (column color bars from outer to inner indicate treatment, tolerance, and subpopulation (I, J); and MAPMAN pathway representation of identified differentially expressed metabolites for treatment*tolerance effects (K, L).

With treatment*tolerance effect, the ind_aus panel had more metabolic features differentially expressed, with 211 up-regulated and 319 down-regulated (Figure 5B). Moreover, the jap_aro panel had 16 up-regulated and 50 down-regulated metabolic features (Figure 5A). Glycerate-3-phosphate substantially decreased (-logfc -1.60**), reflecting its conversion to serine (-logfc 1.10***). TCA is considerably enhanced in tolerant ind_aus genotypes due to more abundant 2-oxoglutarate and fumarate (-logfc of 1.29* and 1.70***, respectively). This is reflected in purported increase in the amino acids glutamate, glutamine, and proline. The plant hormones zeatin, jasmonate and indole-3-acetate significantly declined in tolerant ind_aus genotypes (-logfc of -1.44**, -2.44** and -1.61*, respectively). Lactate substantially increased (-logfc =1.27**) in tolerant ind_aus genotypes implying augmented fermentative metabolism for energy production. Conversely, tolerant jap_aro genotypes up regulate alpha-tocopherol, probably to alleviate ROS injury. Interestingly, cholesterol synthesis decreased in tolerant jap_aro genotypes (-logfc = -1.33**) indicative of intact membrane as a consequence of efficient ROS sequestration. Furthermore, declined lipid synthesis might suggest reduced demand for ATP and O_2_. Bio-unavailable plant hormone forms, particularly glycosylated cytokinins (benzylaminopurine-glucoside, -logfc = 1.60*) declined in tolerant jap_aro genotypes, indicative of dynamic conversion of storage pools to bioactive cytokinin forms.

### Metabolic confluence networks exposes conserved and subpopulation-specific modules

To generate metabolic confluence network, the WGCNA approach was implemented to discover modules of highly correlated metabolic features and relate these expression profiles to its corresponding phenotypic responses, particularly tolerance. Differences in metabolic networks between jap_aro and ind_aus were explored while simultaneously determining biochemical modules and biomarkers contributing to AG tolerance. To assess the global similarities of jap_aro and ind_aus metabolome networks, *kME* values (Eigengene-based network connectivity) for all the metabolic features and their corresponding fluxes were compared. Both network connectivity and fluxes were highly preserved between panels due to strong correlations (0.90*** for kME and 0.98*** for confluences; Suppl. Figure S17). To explore co-expression module changes between metabolomes of the jap_aro and ind_aus genotypes, module preservation was assessed between networks. Results showed high degree of between – subpopulation module preservation, and in fact the blue and turquoise modules of both jap_aro and ind_aus panels significantly overlap (Figure 5C, 5I for jap_aro, 5G, 5J for ind_aus). The blue metabolic module includes compounds involved in carbon and nitrogen pathways while the turquoise metabolic module comprises metabolites from carbon, nitrogen and nucleotide metabolism (Suppl. Dataset S4). The strong module conservation depicts that AG stress dramatically alters the metabolic landscape across subpopulations. The highly correlated metabolites within these modules play a central role in the biochemical repertoire and represent the core metabolic fluxes during germination under low oxygen stress. However, the brown module was found to be jap_aro specific due to very weak module preservation in the ind_aus metabolome (Figure 5D for jap_aro, 5H for ind_aus). The brown module includes identified metabolites: UMP, GMP, alpha-tocopherol, squalene, and vitamin K (Please see Suppl. Datasets S3 and S4 for the module assignment and pathway enrichments for the metabolites).

## DISCUSSION

### Different subpopulations displayed varying AG adaptive responses

Our study had dissected the genetic architecture of AG tolerance in rice and integrated network theory to tease out possible biological processes and molecular functions associated with the response. The phenotypic assessment of the diversity panel revealed the range and distribution of tolerance to AG stress and uncovered insights regarding the evolution of the ability to germinate under flooding conditions among rice subpopulations. Our results indicated prevalence of AG tolerance in japonica subpopulations, though a number of aus and indica representatives showed remarkable tolerance (Figure 1A). Moreover, varying phenotypic responses were deployed by different subpopulations: tolerance is attributed to fast shoot elongation and unhampered root growth for japonicas, but shoot elongation alone for indicas (Figure 1B-C).

Plants had evolved strategies to surpass varying flooding regimes. The escape strategy stimulates rapid elongation of the shoot to gain access to the aerial atmosphere. Though this approach can facilitate gas exchange and photosynthesis, it requires high metabolic demands to sustain growth. The SNORKEL1 and SNORKEL2 genes from C2985 rice cultivar confers tolerance to deep-water flooding through fast elongation as an escape strategy. These genes encode ethylene response factors involved in ethylene signaling, and its expression results in gibberellin-induced internode elongation (<u>Hattori et al. 2009</u>).

Nevertheless, if flooding is ephemeral, this mechanism would be disadvantageous. In such case, quiescent strategy is favored wherein the plant has to restrict growth underwater and conserve resources. As the flooding recedes, the unused reserves can then be utilized for growth and recovery. The *SUB1-A* allele of FR13A rice landrace was found to confer tolerance to complete submergence for 10-14 days through quiescence (<u>Xu et al. 2006</u>). This allele variant encodes an ethylene-response-factor-like transcription factors and its expression results in over accumulation of GA repressors Slender Rice-1 (SLR1) and SLR1 Like-1 concomitant to the brassionosteroid-mediated activation of GA catabolism, consequently abolishing ethylene-promoted- or brassinosteroid-modulated- GA-induced elongation underwater (<u>Fukao and Bailey-Serres, 2008, Schmitz et al., 2013)</u>. For AG stress, the escape strategy is advantageous and the rapid seed carbohydrate mobilization, shoot growth and unhampered root development are required for adaptation despite inefficiency of fermentation under oxygen deprivation. Since the embryo primarily relies on the starchy endosperm, the ability to mobilize the complex carbohydrate to sugar under low-oxygen is critical for germination seedling survival (<u>Ismail et al. 2009</u>).

The negative correlation of starch with growth parameters and survival strongly indicates that tolerant varieties were able to convert starch reserves to simple sugars to address energy crisis and metabolic requirements of the growing embryo (Suppl. Figure S1). Our study attempted to investigate variation in early root growth under AG stress by utilizing root trainers, which permit considerable root growth for the seedling in light of understanding delayed radicle emergence and its eventual protrusion upon coleoptile contact with the water surface. Since the root tissues are most inflicted during oxygen limitations partly due to its proximity to air and also due to associated soil toxicoses from microbial action (<u>Colmer and Voesenek, 2009; Kirk et al., 2014</u>), as root development is equally important with shoot growth under AG stress. Aside from its physiological function, proper soil anchorage could prevent seedling drift and lodging later in the season.

### AG tolerance is associated with numerous genetic loci

To map genomic regions associated with the traits, we have implemented ECMLM algorithm with correction for population stratification to reduce false positive signals (<u>Li et al. 2014</u>). Since results indicate that geographical adaptation and the pattern of natural variation tends to coincide with population structure, analyses were sub-divided within the diversity panel to capture allelic variations segregating within or among subpopulations (Figure 1E). Additionally, AG tolerance being a complex trait, ECMLM resulted in modest associations but with deflated quantile-quantile plots indicative of weak statistical power (Suppl. Figure S2-S5), inappropriateness, or over-correction of the model. Accordingly, new approach was employed and the statistical parameters were substantially improved for most of the traits upon application of the SUPER method. With this, different algorithms should be exploited to evaluate suitability of the model since it is assumed that the genetic landscape underlying natural variation behaves differently across traits (<u>Zhao et al. 2011</u>). Our results were able to detect, to some extent, some of the published QTLs for AG tolerance, at least using this diversity panel; additionally, some of the SNP peaks have not been identified before implying that these regions harbor novel genes or allelic variants as plausible determinants of the phenotypic response. Interestingly, the phenotypic responses were controlled differently among subpopulations, due to varying genetic associations considerably linked with the traits and conspicuous chromosomal hotspots that appear to be segregated among subgroups from the stratified analyses (Figure 2A). These evidences reflect the rich history of rice domestication and suggest disparity in the selection and ecological pressures in the divergence of rice subpopulations. It is also possible that the susceptibility of modern high-yielding varieties to AG stress was due to the dominance of transplanting for crop establishment, wherein farmers had unintentionally selected against plastic germination by providing suitable environment for seedling growth. Similarly, a mutation in the *Waxy* gene responsible for amylose deficiency characterized with non-glutinous phenotype in most temperate japonicas has been found to be subjected to artificial selection. This mutation was absent in other rice subpopulations, and such selective sweep was owed to anthropological influence, particularly cultural practices, and food preferences (<u>Olsen et al. 2006</u>). In rice, the *Sdr4* gene was cloned and found to be involved in the regulation of seed dormancy (<u>Sugimoto et al. 2010</u>). Domesticated crops like rice, has been artificially selected to obtain rapid and synchronous germination for successful cultivation. Conversely, seed dormancy is typical for wild species as it confers protection to harsh environments, prevents competition within species, and avoids out-of-season germination (<u>Finkelstein et al. 2008</u>). The japonica cultivar Nipponbare had the *Sdr4-n* allele which causes them to lose dormancy immediately resulting on pre-harvest sprouting but, the indica cultivar Kasalath had *Sdr4-k* allele conferring considerable dormancy (<u>Sugimoto et al. 2010</u>). These subpopulation differences reflect the divergent domestication routes for indica and japonicas, which may implicate their disparity in dormancy, germination in anaerobic soils and its plasticity. Furthermore, these evidences seemingly suggest that certain domestication traits are preferentially sorted among subpopulation.

It is also noteworthy that wild relatives of rice could be utilized as novel sources of tolerance genes, which have probably undergone domestication sweep through the evolutionary course, to facilitate breeding efforts (<u>Tanksley and McCouch, 1997</u>). Traits with significant associations include shoot- and root-related traits, and starch, which concurred with many other traits implicating pleiotropic effects of the associated genomic region (Figure 2A). Apparently, it is more perceptive to dissect complex traits to component traits to increase detection power of association mapping (<u>Crowell et al, 2016</u>). This approach could facilitate teasing-out genetic control of complex traits into numerous loci of considerable signals. Due to crossability issues between indica and japonicas, alleles for AG tolerance within subpopulation pools can offer breeding convenience. However, pyramiding different genetic variants across subpopulations controlling the trait proposes possibilities of conferring robust AG tolerance for varietal improvement, especially that subpopulations offer varying adaptive responses. Yet some of these genetic regions exhibited very low heritability estimates and occur at low frequencies, which are expected for traits that are inherently complex, overlap with population structure and highly influenced by environment (<u>Brachi et al. 2011</u>). It is possible that the low heritability is attributed to epistasis or the inherent allelic architecture of the trait: common variants with very small effects, rare variants with large effects, and structural or copy number variants that could not be captured by SNP arrays (<u>Manolio et al. 2009</u>; <u>Ingvarsson and Street. 2011</u>). This study has demonstrated the expanse of natural variations for AG tolerance, the complexity of its genetic control, and evolutionary glimpses of the trait existing in the rice diversity panel. The tolerant genotypes identified here will be crossed to develop populations for QTL studies, further genetic validation, and breeding platforms to facilitate crop improvement for direct seeded systems.

### Integration of network analysis and functional enrichment complements and strengthens GWA mapping

Though GWAS has been successful and increasingly discovering genomic regions significantly associated with traits of interest, genetic variants of small effects but with true association suffer from stringent statistical thresholds (Figure 2, Suppl. Figure S11). Likewise, abstraction of biological annotations from these genomic scans has recently gained greater attention (<u>Wang et al. 2010</u>; <u>Peng et al. 2010</u>). To address these limitations, complementary approaches, specifically gene network analyses coupled with pathway-based enrichment have been utilized. Our study made use of publicly available microarray data of similar experiments to construct gene networks and group the candidate genes in highly correlated modules that are condensed for specific mechanisms and functions. With this, we have identified modules of high biological pertinence to the AG response on the bases of network metrics (Figure 3 and Suppl. Figures S9-S10). Module characterization had revealed specific biological processes and molecular functions that have not been uncovered by conventional pathway-based analysis of the entire gene list (Suppl. Dataset S2).

Module enrichments have shown the fundamental involvement of carbohydrate metabolism in AG tolerance due to significant involvement of many modules with primary metabolism, carbohydrate transport and sugar homeostasis. It has been established that, under oxygen limitation, the conversion of starchy endosperm to simple sugar is critical to address ATP crisis and precursor demands for the growing embryonic axis (<u>Guglielminetti et al. 1995</u>; <u>Perata et al. 1997</u>; <u>Hwang et al. 1999</u>). Studies had also shown that sugars serve as a signaling molecules under oxygen deprivation. The CIPK15 protein integrates sugar- and O_2_-deficiency signaling by mediating the SNRK1-MYBS1-mediated-sugar sensing cascade to modulate carbohydrate catabolism and fermentation, thus leading to seed germination and seedling survival when flooding occurs following direct seeding (<u>Lee et al. 2009</u>). Recently, the genetic determinant OsTPP7 derived from KHO was identified, which modulates sugar signaling through reduction of T6P pools, consequently increasing sucrose for growth of the embryo in rice germinating under flooded conditions (<u>Kretzschmar et al. 2015</u>). Artificial application of T6P analogues enhances grain yield and augments performance under stress conditions (<u>Griffiths et al, 2016</u>).

Some of the modules had functional enrichments for biological regulation at levels of transcription and translation revealing extensive effect of oxygen deprivation in to the global gene expression (Suppl. Dataset S2). In *Arabidopsis*, hypoxia stress extensively altered translatomes in which selective translation purports less energy consumption. Hypoxia-induced translatome modification reflected substantial changes in transcription, mRNA turnover and translation. Under anoxia, most of the ATP synthesized was spent for translation in rice coleoptiles (<u>Edwards et al, 2012</u>). On the other hand, transcriptome landscape under hypoxia showed remarkable decline in ribosome loading with mRNAs as a repercussion of translational prioritization to conserve energy (<u>Mustroph et al. 2009</u>; <u>Branco-Price et</u> <u>al. 2005</u>). Moreover, recent evidences implicate importance of translational control via protein degradation under O_2_ deprivation and were found to be integral in metabolic economy and signaling cascades under hypoxia. The transcription factor family VII ETHYLENE RESPONSE FACTORS (ERF-VII), which includes *SUB1* and *SNK1* were established to be the modulator of hypoxia-responsive genes. Its protein stability is controlled in an O_2_-dependent fashion through the N-end rule pathway of targeted proteasomal degradation (<u>Gibbs et al 2011</u>; <u>Licausi et al, 2011</u>).

Many of the modules interestingly uncover participation in signaling cascades including MAPK cascade, G-protein receptor signaling and, serine/threonine kinase signaling pathway, which are implicated in many biological processes and environmental responses (Suppl. Dataset S2). Recently, MAPK cascades that modulate tolerance to complete submergence were reported in rice. The MPK3 protein selectively phosphorylates the tolerant *SUB1-A1* allele; conversely *SUB1A-1* interacts with the promoter of MPK3, ensuing a positive feedback regulation (<u>Singh and Sinha, 2016</u>). In Arabidopsis, the MAPK phosphatase PP2C5, and attenuates MAPK activation, adversely affecting several responses under stress conditions, including stomatal opening, seed germination, and ABA-regulated gene expression (<u>Brock et al. 2010</u>).

Similarly, the serine/threonine protein kinase TaSnRK2.4 in wheat enhances tolerance to drought, salt, and cold stress (<u>Mao et al. 2009</u>). Similarly, G proteins were found to promote lysigenous aerenchyma formation via programmed cell death under submergence, ethylene, and H_2_O_2_ treatment in rice (<u>Steffens and Sauter, 2009</u>). Our study had liberated specific biological processes and molecular functions involved in AG tolerance by integrating GWA mapping with gene network co-expression analysis and pathway-based enrichment characterization.

### Incorporation of network metrics facilitates pragmatic candidate gene nomination

Network topology, aside from GWAS p-values, can be used as an impartial parameter for selecting candidate genes. Using this metric offers advantage of reproducibility, though further validation is still needed (<u>Farber, 2013</u>). Here, we have demonstrated the use of the network index GS to objectively select plausible genetic determinants of the trait among stratified analyses (Figure 3, and Suppl. Figures S9-S10). This network index reflects the correlation of the phenotypic response with the expression profile of a particular gene (<u>Langfelder and Horvath, 2008</u>). Our results uncovered functional connections of the candidate genes with highly connected neighboring genes within the module, further teasing out biological mechanisms at the subnetwork level (Suppl. Figure S11). This pipeline can be exploited in cases of proteins of unknown functions that seemed to be putative genetic determinants, in which inspection of the submodule functions could lead to inferred roles of the unclassified protein. These inferences could also provide information regarding regulation of the candidate genes by identifying proteins that interact or were co-regulated with it. Network analysis could also identify hub genes, which had extensive influence in the entire expression network thus had a key role in the biology of the response. With this approach we were able to empirically select candidate genes using systems-level approach, add value through network biological deduction and identify allelic variants conferring AG tolerance (Suppl. Figures S12-S14). The identified discriminative SNP markers can further be used for QTL analysis and marker-assisted breeding. The identified genes still warrants further validation through population study, reverse genetics or genome editing.

### Metabolic signatures and module preservation across subpopulations define core and subpopulation-specific shifts under AG

To assess the metabolic landscape under AG perturbation with the aim of understanding metabolic shifts as an adaptive response to oxygen limitation, systems-biology approach was implemented. AG causes intensive and extensive alteration in the metabolome across subpopulations, particularly the central carbon and nitrogen metabolism (Figure 5, Suppl. Datasets S3 and S4). This suggests that processes involved in seed germination are highly conserved and tightly controlled. Similarly, metabolomes of seeds and its development showed highly synchronized pattern indicative of tightly regulated and highly conserved regulation among indica and japonica rice cultivars (<u>Hu et al, 2014</u>; <u>Hu et</u> <u>al, 2016</u>). Metabolic adjustments under anoxia include increased glycolytic activity and induced fermentative pathways (<u>Bailey-Serres and Voesenek, 2008</u>) to compromise energy demands and conserve and recycle precursors. Under anoxic conditions, carbohydrate metabolism is extremely dampened primarily through inhibited starch degradation and secondly by impeded pyruvate entry into the TCA cycle (<u>Miro and Ismail, 2013</u>). Our results further validate these adjustments, in particular the increased abundance of sugars feeding the glycolysis; the accumulation of pyruvate and its fermentative products lactate and alanine due to inhibited TCA; and the subsequent fluctuations in amino acid levels (Figure 5K-L). However there are certain degrees of differences among subpopulations, reflecting specific adaptations that correspond with divergent domestication routes and these metabolic variations may partly explain dissimilarities in AG tolerance strategies (Figure 5C-D, 5G-H). A small but distinct biochemical network was found to be specific for jap_aro panel with definite involvement in ROS alleviation, signaling and lipid metabolism (Figure 5D, Suppl. Datasets S3 and S4). The inherent metabolic variations occurring in seeds among subpopulation may have eventual effects on stress responses, particularly in breaking dormancy and during the germination phase (<u>Hu et al, 2013</u>). Though jap_aro generally had shown better TCA metabolism, the tolerant ind_aus genotypes had fairly promoted TCA cycle to efficiently recycle and better reconfigure resources; consequently, providing more and diverse amino acids. Conversely, tolerant jap_aro genotypes had improved ROS sequestration via antioxidants and efficient lipid synthesis, which should result in improved membrane fluidity (Figure 5D, Suppl. Datasets S3 and S4). Lipid metabolism is important for maintaining membrane integrity and may have substantial repercussion in sensing and signaling pathways in plants (<u>Edwards et al, 2012</u>; <u>Penfield, 2008</u>). Metabolic profiles of subpopulations are accompanied by changes in plant hormones that fluctuate between ind_aus and jap_aro panels. These changes suggest that subpopulations might deploy varying signaling machineries in regulating germination under unfavorable conditions.

## CONCLUSIONS

This study dissected the underlying genetics of tolerance to flooding during germination in rice, unveiling varying genetic controls in concordance with geographical adaptation and subpopulation structure, typical of traits that are complex in nature. Different subpopulations confer varying AG escape strategy, particularly involving preferential tissue growth with corresponding metabolic adjustments. AG stress resulted in dramatic shifts in the metabolomes across subpopulations comprising core biochemical fluxes involved in central metabolism. However a small community of metabolic repertoire differentiates the subpopulations, which play integral role in ROS sequestration, signaling, and lipid synthesis. Integration of transcriptome information through network theory strengthens GWA mapping by providing insightful information on the functional processes involving gene modules of high relevance to the trait and offers empirical bases of nominating genes for further validation. Allelic variants of selected candidate genes were identified and strongly correlate with the phenotype. The discriminative SNPs identified can be utilized as a diagnostic tool to facilitate MABC and breeding efforts. Populations will be developed from selected tolerant genotypes for QTL analysis to expedite breeding for different rice ecologies. This study provided a comprehensive systems-level approach for deciphering the genetics and physiology of AG tolerance by consolidating genetic architecture, transcriptomic meta-analysis and, metabolite profiling in a global and genome-wide perspective.

## MATERIALS AND METHODS

### Plant materials and growth conditions

The experiments were conducted in greenhouses at the International Rice Research Institute using a rice diversity panel consisted of 343 accessions: 53 admixed (adm), 12 aromatics (aro), 56 aus, 72 indica (ind), 74 temperate japonica (tej), and 76 tropical japonica (trj) (Table S1). The panel was screened for AG tolerance, with Mazhan Red (tolerant) and IR42 (sensitive) used as checks. Prior to sowing, seeds were heat treated at 50 °C for 5 days to break dormancy. The first experiment used seed trays in randomized complete block design with three replications. Twenty seeds per genotype were sown in seed trays half-filled with soil, then covered with approximately 1 cm soil. One set was maintained in water-saturated soil as control; the second set was submerged under 10-cm water depth immediately after dry seeding and maintained for 21 days. Seedling emergence was assessed at 14 and 21 days after sowing. To investigate root traits, the second experiment utilized Deep Rootrainers^TM^ (Tildenet), comprising of 32 compartments per tray, with dimensions of 36 cm x 21 cm x 12 cm, holding 175 cm^3^ soil to permit largely undisturbed root growth; was done in randomized complete block design with three replications. Three seeds were sown in each of four compartments per genotype, then covered with 1 cm soil. One set was saturated with water as control while the other was subjected to AG stress. Survival was assessed 14 days after sowing.

For metabolite profiling, surface-sterilized seeds of genotypes selected based on their contrasting responses to AG stress across subpopulations were placed in 0.75% agar with 0.5 strength Yoshida solution in a test tube and poured with approximately 1 cm of same medium with 0.10% activated carbon to prevent light penetration mimicking the soil. The agar portion was covered with aluminum foil to avoid light exposure. The AG stress was imposed by aseptically pouring in autoclaved water to a depth of 10 cm from the agar surface. The setup was placed in growth chambers (CONVIRON) set with 12:12 photoperiod at temperatures of 28C/25C (day/night) and 350 µE m^-2^ s^-1^ light intensity.

### Phenotyping of rice accessions

Number of seedlings that emerged over water was scored as percent (%) emergence at 14 and 21 DAS. Likewise, the number of seeds that were able to emerge but stayed under water was also noted as % germination. The emergence and germination index was computed by dividing the % emergence and % germination with the number of seedlings germinated under control conditions. The lengths of the shoot, leaf sheath, and leaf blade were also measured. Shoot area at 21 DAS was estimated from pictures through ImageJ software (<u>Schneideer et al. 2012</u>). The dry weights of the shoots and roots were gathered by drying the dissected shoots and roots at 21 DAS for 5 d at 70 °C. Root morphological traits at 14 DAS were assessed using scanning coupled with analysis through WinRhizo root imaging analysis software (<u>Arsenault et al. 1995</u>).

Chlorophyll and carotenoids were determined spectrophotometrically following <u>Lichtenthaler and</u> <u>Buschmann (2001)</u>, with modifications. Leaf samples were lyophilized and extracted with 95% ethanol for 24 hours at 25°C. Absorbance was recorded at 470, 645, and 664nm and pigment concentrations were calculated on a dry weight (DW) basis. Ethanol-soluble sugar in seeds was assayed according to <u>Fales (1951)</u> with modifications. Freeze-dried samples were finely ground and extracted with 80% ethanol. The extract was reacted with 7.5 mM anthrone reagent and absorbance read at 620 nm. Sugar concentrations were then determined using dextrose standard curve. The final sugar content was expressed on DW basis. The resulting residue was subjected to starch assay based on <u>Kunst et al. (1988)</u>, with slight adjustments. The residue was digested with 25 mM Na acetate buffer (pH 4.6), and 15 units amyloglucosidase enzyme (Sigma Aldrich). The hydrolysate was reacted with PGO-enzyme: 1 unit peroxidase and 5 unit glucose oxidase (Sigma Aldrich), and 0.12 mM o-dianosidine dihydrochloride (Sigma Aldrich) for 30 min in the dark and absorbance was read at 450 nm. Starch concentration was determined through soluble starch standard curve and final content was expressed on DW basis (Suppl. Table S2) for the list of measured and derived traits).

### Statistical Analysis

Phenotypic data was inspected through histogram, quantile-quantile plot, and residual plot; and Box-Cox transformed when necessary to follow the linear model assumptions: independence, normality, and homoscedasticity, prior to analysis. Analysis of variance with Tukey’s HSD for post-hoc test, Pearson correlation, and multivariate analysis were accomplished using R and STAR software (<u>R Core Team,</u> <u>2013</u>; <u>IRRI, 2013</u>). Transcriptomic and metabolomics data had undergone Empirical Bayesian inference, correlation, and multivariate analysis for statistical evaluation.

### Genome-wide association mapping

The mean value of three biological replicates was used as the phenotype data for association mapping. Phenotypic data structure was examined through residual analysis and assumed to fit Gaussian distribution. If necessary, the data has undergone Box-Cox transformation (<u>Box and Cox, 1964</u>), to follow the assumptions of the linear model prior to analysis. Genome wide association mapping was conducted with the transformed phenotypic data and the 44K SNP dataset (<u>Zhao et al. 2011</u>) using GAPIT in R (<u>Tang</u> <u>et al. 2016</u>). The SNP array sufficiently provides genomic resolution of one SNP every 10 kb for the entire 12 rice chromosomes. SNPs with minor allele frequency (MAF) of < 10% were filtered out, leaving 32,175 SNPs used for association. An Enriched Compressed Mixed Linear Model (ECMLM) and Settlement of Mixed Linear Model Under Progressively Exclusive Relationship (SUPER) method were implemented, accounting also for population stratification by including the first three principal components and kinship matrix to the model (<u>Li et al. 2014</u>; <u>Wang et al. 2014</u>). Association mapping was conducted for the entire panel and between subpopulations.

### Generation of candidate gene lists

The top 20 SNP peaks for every trait were further inspected to generate candidate gene lists. Significant SNPs within 200kb region (the mean LD decay across rice subpopulations) from each side with the SNP as the center, were merged to constitute LD blocks, these resulting regions (flanked with 100kb, either as LD blocks or singleton) were examined in the Gramene genome browser for genes and considered as putative determinants for the trait (<u>Tello-Ruiz et al. 2016</u>).

### Gene Ontology and pathway –enrichment analysis

The candidate gene lists and network modules were analyzed for statistically over- and under- represented functions using the Protein ANalysis THrough Evolutionary Relationships database (<u>Mi et al.</u> <u>2013</u>; <u>Gene Ontology Consortium, 2013</u>). The gene list is compared with the reference gene list (the set of all genes of *Oryza sativa*) through binomial distribution testing for each of the Gene Ontology, Panther protein class, or pathway terms.

### Gene expression data processing

To generate gene co-expression networks, published microarray data from relevant experiments were used (<u>Lasanthi-Kudahettige et al. 2007</u>; <u>Howell et al. 2007</u>; <u>Narsai et al. 2009</u>). These studies involved transcript profiling of germinating rice seeds under normoxic (8 samples) and anoxic (8 samples) conditions, with corresponding coleoptile lengths imputed. The Affymetrix CEL files were downloaded from the Gene Expression Omnibus of National Center for Biotechnology Information (Dataset GSE6908, https://www.ncbi.nlm.nih.gov/geo/query/acc.cgi?acc=GSE6908) and from ArrayExpress of European Molecular Biology Laboratory – European Bioinformatics Institute (Dataset E-MEXP-2267, http://www.ebi.ac.uk/arrayexpress/experiments/E-MEXP-2267/). The raw data were processed using the *affy* package (<u>Gautier et al. 2004</u>) for the R Language and Environment for Statistical Computing (<u>R</u> <u>Core Team, 2013</u>). Robust multiarray algorithm was implemented to extract normalized probe level expression data from the samples (<u>Irizzary et al. 2003</u>). The expression profiles were corrected for batch effects using the *sva* package (<u>Leek et al. 2012</u>).

### Transcriptomic network analysis

Network analysis was performed using WGCNA (weighted gene co-expression network analysis) R package (<u>Langfelder and Horvath, 2008</u>). An extensive guideline for WGCNA, including tutorials are found at http://www.genetics.ucla.edu/labs/horvath/CoexpressionNetwork/. A WGCNA network for the selected probes was constructed with manual module. The package initially implements a soft threshold as fitting index to assess scale-free network constructed upon gene-gene correlation from microarray samples. Genes were clustered based on topological overlap of their connectivity through average linkage hierarchical clustering followed by dynamic tree cutting to define network modules. Gene Significance (GS) for each gene network was defined as the Pearson correlation with trait while Module Significance (MS) is the mean of absolute GS values of the genes in the module. Module Membership (MM) was computed as the Pearson correlation between each gene’s expression and its module eigengene. Topological parameters from the network were calculated using NetworkAnalyzer (<u>Doncheva et al. 2012</u>) and the network depictions were visualized using Cytoscape (<u>Shannon et al.</u> <u>2003</u>).

### Haplotype analysis

Candidate genes from the selected modules of strong biological relevance were selected on the basis of high absolute GS values (≥ 0.80) and its associated regions (SNP singleton or LD blocks with –log10 p- values > 5.81) were analyzed for haplotypes using Haploview (<u>Barrett et al. 2005</u>). The haplotype blocks of strong linkage disequilibrium (LD) were further inspected for connection with AG tolerance. Subnetworks of the selected candidate genes were also depicted using qgraph in R (<u>Epskamp et al.</u> <u>2012</u>) to reveal highly correlated neighboring genes and possible pathways enriched (<u>Mi et al. 2013</u>). In- silico analysis was also extended by utilizing SNPs from the database of SNP- seek providing SNP calls for 3000 genomes (http://snp-seek.irri.org/, <u>Mansueto et al, 2017</u>).

### Non-targeted metabolite profiling of polar to semi-polar fractions

Germinating seeds were sampled at 4 DAS and temporarily stored at -80C. Samples were extracted with 0.3:0.79:0.01:2.5 methanol: isopropanol: glacial acetic acid: water spiked with internal standards (ribitol, norleucine 10 μg mL^-1^ each) and chloroform spiked with internal standard (5-α-cholestane, 10 μg mL^-1^). Polar fractions were lyophilized and derivitised with methoxamine in pyridine (20 mg mL^-1^) and MSTFA spiked with retention index markers C7-C36 alkanes (1000 µg mL^-1^). Samples were injected in Shimadzu GCMS-QP2010 Plus with SLB 5MS column, 1µL split-less with initial temperature of 60°C ramped at 5°C min^-1^ till 330°C. Full scan mass spectra were recorded through a range of 40-600 m/z with scan rate of 1250AMU s^-1^ (<u>Lisec et al, 2006</u>).

### Chromatogram pre-processing

The TargetSearch pipeline for preprocessing of GC-MS data was implemented in R (<u>Cuadros-Inostroza et</u> <u>al, 2009</u>). The CDF files were baseline corrected and normalized based on day measurements and peaks were identified, adjusted for retention time, and evaluated for outliers. The peaks were matched against the reference library from Golm Metabolome Database (<u>Hummel et al, 2013</u>) and the metabolite profile was generated. The intensities of metabolites were normalized with their corresponding internal standards then with fresh weight sample-wise. The missing values of a given metabolite were imputed with the detected minimum for statistical analysis, with the assumption that it is below detection limits.

### Metabolic Network construction and module preservation calculation

WGCNA pipeline was utilized to construct co-expression network, identify modules, and determine module preservation from the metabolite profile of the ind_aus and jap_aro consisting of 40 and 48 samples, respectively, with 1355 detected metabolic features (<u>Langfelder and Horvath, 2008</u>; <u>Langfelder</u> <u>et al, 2011</u>). The absolute Pearson correlation coefficients were computed for pair-wise comparison of metabolites across samples. Outliers were removed and the resulting matrix was tested for scale free topology approximation. The dataset lacked scale free topology and the heterogeneity is attributable to strong treatment effect resulting in high correlations among metabolites, however the mean connectivity remained considerably high. Adjacency matrix was derived from the dataset using a soft power of 16 (for 30-40 samples) and computed dissimilarity based on topological overlap to construct networks. The metabolic data were also associated with the AG tolerance to identify highly pertinent metabolites and modules. The module preservation statistics were computed accordingly with the jap_aro metabolic network as the reference network and the ind_aus metabolic network as the test. Median rank was used to detect module preservation and Zsummary to evaluate significance of module preservation with 200 permutations. Modules with Zsummary score > 10 indicates strong preservation, below 10 to 2 indicate weak to moderate preservation and < 2 indicate no preservation. Topological parameters from the networks were calculated using NetworkAnalyzer (<u>Doncheva et al. 2012</u>) and the network depictions were visualized using Cytoscape (<u>Shannon et al. 2003</u>).

### Metabolic pathway-enrichment analysis

The resulting metabolic modules were analyzed for over- representation of pathways using web-based IMPaLA database (http://impala.molgen.mpg.de/, <u>Kamburov et al, 2011</u>). The database contained 3073 annotated pathways from 11 public databases including KEGG (http://www.genome.jp/kegg/), Reactome (http://www.reactome.org) and, Wikipathways (http://www.wikipathways.org). Based on the list of the identified metabolites from each module, hypergeometric distribution is utilized to generate significance values for each pathway in terms of the overlap with those associated with the list.

## ACKNOWLEDGMENTS

We thank the Genetic Resources Center of IRRI for providing the seeds; Berta Miro for giving useful insights in the analysis; Shaira Gapuz, Ofelia Namuco, and Melencio Apostol for technical assistance in the experiment. This work was supported by the Stress Tolerant Rice for Africa and South Asia (STRASA) project funded by the Bill and Melinda Gates Foundation (BMGF) and a grant from the German Federal Ministry of Economic Cooperation and Development (BMZ) #81157485 to A.M.I.

## COMPETING FINANCIAL INTERESTS

The authors declare no competing financial interests.

## SUPPLEMENTARY MATERIALS

**Supplementary Table S1**. The accessions included in the Rice Diversity Panel 1 with their corresponding subpopulations and geographical origin.

**Supplementary Table S2**. The list of measured and derived traits from seedling tray and root trainer screening methods.

**Supplementary Table S3**. Significant genetic loci (-log10 p-values > 5.81, Bonferroni cut-off) associated with AG traits from stratified GWA mapping.

**Supplementary Figure S1**. Correlation of measured and derived phenotypic traits. The side color scale bar indicates Pearson coefficients.

**Supplementary Figure S2**. GWAS results for measured and derived traits of the all panel as depicted by Manhattan plots, MAF graphs, quantile-quantile plots and, heritability estimates using ECMLM algorithm.

**Supplementary Figure S3**. GWAS results for measured and derived traits of the ind_aus panel as depicted by Manhattan plots, MAF graphs, quantile-quantile plots and, heritability estimates using ECMLM algorithm.

**Supplementary Figure S4**. GWAS results for measured and derived traits of the jap_aro panel as depicted by Manhattan plots, MAF graphs, quantile-quantile plots and, heritability estimates using ECMLM algorithm.

**Supplementary Figure S5**. GWAS results for measured and derived traits of the all panel as depicted by Manhattan plots and, quantile-quantile plots using SUPER algorithm.

**Supplementary Figure S6**. GWAS results for measured and derived traits of the ind_aus panel as depicted by Manhattan plots and, quantile-quantile plots using SUPER algorithm

**Supplementary Figure S7**. GWAS results for measured and derived traits of the jap_aro panel as depicted by Manhattan plots and, quantile-quantile plots using SUPER algorithm.

**Supplementary Figure S8**. MAF distribution and plots of MAF against significance values and number of associated traits for all (A), ind_aus (B) and, jap_aro (C) panels.

**Supplementary Figure S9**. Venn diagram of differentially expressed genes for ind_aus panel (A). Co- expression network from the ind_aus panel, each line is an individual gene and the branches correspond to modules of highly interconnected genes, below the dendogram, each gene is color coded to indicate module assignment and the GS values for coleoptile length (B). Correlation between MM and GS for each of the identified GWAS modules (C).

**Supplementary Figure S10**. Venn diagram of differentially expressed genes for jap_aro panel (A). Co- expression network from the jap_aro panel, each line is an individual gene and the branches correspond to modules of highly interconnected genes, below the dendogram, each gene is color coded to indicate module assignment and the GS values for coleoptile length (B). Correlation between MM and GS for each of the identified GWAS modules (C).

**Supplementary Figure S11**. GWAS and gene co-expression plots for all (A), ind_aus(B) and, jap_aro (C) panels; first track includes published QTLs linked with AG tolerance derived from different donors, dark red (KHO), dark green (Nanhi), black (Mazhan Red); second track indicates position of top 20 SNPs from each trait, with the bar representing –log10 p-values, filled circles identify significant associations; third track reflects minor allele frequency from ECMLM approach; fourth track shows number of traits associated with the SNPs; the center depicts the interaction of the tightly correlated genes (GS ≥ 0.80 with the 97.5 percentile of the interactions plotted), color indicate the module assignment upon WGCNA implementation.

**Supplementary Figure S12**. Ideogram depiction of genes present in ind_aus LDb09010 region in chromosome 9 with corresponding GWAS significance values and MAF (A). Linkage disequilibrium heatmap of the region showing 4 haplotype blocks (B). Haplotype variants existing in the associated region with corresponding linkage disequilibrium values between blocks and the frequencies of the variants (C). Frequency distribution of the identified allelic variants and corresponding tolerance (D).

**Supplementary Figure S13**. Ideogram depiction of genes present in jap_aro id3013397 SNP in chromosome 3 with corresponding GWAS significance values and MAF (A). Linkage disequilibrium heatmap of the region showing no define haplotype blocks (B). Frequency distribution of the identified allelic variants and corresponding tolerance (C).

**Supplementary Figure S14**. Ideogram depiction of the nominated gene LOC_Os08g34580 and the MAF of SNP variants extracted from the 3000 genomes (A). Hierarchical clustering of the genomic region with cutree =12, bar indicates time scale of evolution (B). The overlap of 3000 genomes and the Rice Diversity Panel resulted to 15 genotypes and clades and percent mismatch for Nipponbare genome were plotted with tolerance (C).

**Supplementary Figure S15**. Ideogram depiction of the nominated gene LOC_Os09g26310 and the MAF of SNP variants extracted from the 3000 genomes (A). Hierarchical clustering of the genomic region with cutree =12, bar indicates time scale of evolution (B). The overlap of 3000 genomes and the Rice Diversity Panel resulted in 15 genotypes and clades and percent mismatch for Nipponbare genome were plotted with tolerance (C).

**Supplementary Figure S16**. Ideogram depiction of the nominated gene LOC_Os03g50220 and the MAF of SNP variants extracted from the 3000 genomes (A). Hierarchical clustering of the genomic region with cutree =12, bar indicates time scale of evolution (B). The overlap of 3000 genomes and the Rice Diversity Panel resulted to 15 genotypes and clades and percent mismatch for Nipponbare genome were plotted with tolerance (C).

**Supplementary Figure S17**. Correlation plots of jap_aro metabolic flux with ind_aus metabolic flux (A) and the computed Eigengene-based network connectivity of jap_aro metabolites and ind_aus metabolites (B).

**Supplementary Figure S18**. Gene coexpression network depictions with module size and GS distribution in boxplot, dotted horizontal line indicate module significance, MS ; and histogram of connection strengths, vertical dotted line indicate mean, solid vertical line indicate median, for the modules of the all panel: blue (A), brown (B), green (C), red (D), turquoise (E) and, yellow (F).

**Supplementary Figure S19**. Gene coexpression network depictions with module size and GS distribution in boxplot, dotted horizontal line indicate module significance, MS ; and histogram of connection strengths, vertical dotted line indicate mean, solid vertical line indicate median, for the modules of the ind_aus panel: blue (A), brown (B), turquoise (C), yellow (D), and green (E).

**Supplementary Figure S20**. Gene coexpression network depictions with module size and GS distribution in boxplot, dotted horizontal line indicates module significance, MS ; and histogram of connection strengths, vertical dotted line indicates mean, solid vertical line indicates median, for the modules of the jap_aro panel: blue (A), brown (B),green (C), red (D), turquoise (E) and, yellow (F).

**Supplementary Figure S21**. Correlation between MM and GS for each of the identified metabolic modules in the jap_aro metabolome, all of the modules had significant correlations (p-values < 0.01) for GS and MM.

**Supplementary Figure S22**. Correlation between MM and GS for each of the identified metabolic modules in the ind_aus metabolome, all of the modules had significant correlations (p-values < 0.01) for GS and MM.

**Supplementary Dataset S1**. Complete list of GWAS-WGCNA gene module assignments for all, jap_aro, and ind_aus panels.

**Supplementary Dataset S2**. Complete list of the pathway enrichments of the GWAS-WGCNA gene modules for all, jap_aro, and ind_aus panels.

**Supplementary Dataset S3**. Complete list of the metabolite module assignments for jap_aro and ind_aus metabolomes.

**Supplementary Dataset S4**. Complete list of the pathway enrichments for metabolite module assignments for jap_aro and ind_aus metabolomes.

